# Amplification and attenuation of noisy expression by export processes

**DOI:** 10.1101/2021.10.06.463423

**Authors:** Madeline Smith, Mohammad Soltani, Rahul Kulkarni, Abhyudai Singh

**Affiliations:** Department of Electrical and Computer Engineering, University of Delaware, Newark, DE USA 19716.; Department of Physics, University of Massachusetts Boston, Boston, MA USA 02125.; Department of Electrical and Computer Engineering, Biomedical Engineering, Mathematical Sciences, Center for Bioinformatics and Computational Biology, University of Delaware, Newark, DE USA 19716.

## Abstract

Inside mammalian cells, single genes are known to be transcribed in stochastic bursts leading to the synthesis of nuclear RNAs that are subsequently exported to the cytoplasm to create mRNAs. We systematically characterize the role of export processes in shaping the extent of random fluctuations (i.e. noise) in the mRNA level of a given gene. Using the method of Partitioning of Poisson arrivals, we derive an exact analytical expression for the noise in mRNA level assuming that the nuclear retention time of each RNA is an independent and identically distributed random variable following an arbitrary distribution. These results confirm recent experimental/theoretical findings that decreasing the nuclear export rate buffers the noise in mRNA level, and counterintuitively, decreasing the noise in the nuclear retention time enhances the noise in the mRNA level. Next, we further generalize the model to consider a dynamic extrinsic disturbance that affects the nuclear-to-cytoplasm export. Our results show that noise in the mRNA level varies non-monotonically with the disturbance timescale. More specifically, high- and low-frequency external disturbances have little impact on the mRNA noise level, while noise is amplified at intermediate frequencies. In summary, our results systematically uncover how the coupling of bursty transcription with nuclear export can both attenuate or amplify noise in mRNA levels depending on the nuclear retention time distribution and the presence of extrinsic fluctuations.

## 1 Introduction

Gene expression is the process by which a cell converts information stored within its DNA into functional gene products. The level of gene product has been found to exhibit considerable variation within genetically identical cells of the same population. This is referred to as gene expression noise [1, 2]. Variability can cause deleterious effects within the cell. For example, housekeeping genes encode types of proteins that carry out necessary cellular functions. The prevalence of noise can be detrimental as the level of these essential proteins must be tightly maintained to ensure proper functioning [3–5]. The study of gene expression noise and the mechanisms that drive it are an immense area of interest [6–11].

Gene expression noise is driven by intrinsic and extrinsic sources [12–15]. We consider intrinsic noise as variation that arises due to the inherently stochastic biochemical reactions involved in gene expression, such as transcription and translation. The key players involved in these reactions, like RNA and mRNA, exist at low copy numbers, thus amplifying variation. Moreover, extrinsic noise is the variation due to cell-to-cell factors, such as fluctuations in the cell environment, cell growth, and cell cycle stage [16–19]. In this analysis, we first consider a standard gene expression model featuring intrinsic variations arising from bursty expression, then consider effects on overall noise in the presence of an extrinsic disturbance.

Within the events of gene expression, we are specifically interested in the RNA export process. RNA is a molecule that exports genetic information from DNA stored in the cell nucleus into the cytoplasm as mRNA; a key step in gene expression. The RNA is transported through nuclear pore complexes mediated by nuclear export receptors [20]. Studies reveal that variations in export, such as changes in pathway machinery or export factors, can lead to disease [21]. Therefore, we aim to provide insight into the role of export processes in shaping downstream gene expression noise.

The contribution of nuclear export processes in modulating stochastic gene expression is studied by considering how noise is affected when the (1) export timing is varied, (2) export process includes a time delay, and (3) export occurs in the presence of an extrinsic disturbance. The paper is structured as follows. In section II, we introduce the gene expression model formulation. In section III, we use Partitioning of Poisson Arrivals (PPA) to obtain a general formula for describing noise in cytoplasmic mRNA in the absence of extrinsic noise. The formula is used to characterize the effects on mRNA noise for different RNA nuclear retention time distributions. We further present results when nuclear RNA export includes extrinsic disturbances. Finally, conclusions are presented in section IV.

## 2 Model Formulation

We introduce a stochastic gene expression model with variability due to intrinsic sources. In this setup, gene expression begins in the cell nucleus where we consider a constitutive gene (i.e., the gene is always transcriptionally active) with no feedback regulation. The gene promoter produces bursts of nuclear RNA transcripts, modeled as a bursty birth-death process [22–28]. The bursting events occur at a rate *k*_*x*_ as per a Poisson process to create *B* number of RNA transcripts, where *B* is a discrete random variable drawn from the probability distribution

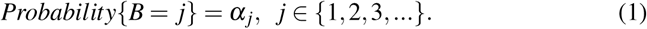

RNA transcripts subsequently undergo export from the nucleus into the cytoplasm. Export of a single RNA molecule occurs at rate *t*_*x*_ to create a messenger RNA (mRNA) transcript in the cytoplasm. The mRNA decays with rate *γ*_*x*_. Prior work has shown that in the limit of rapid export (*t*_*x*_ → ∞) the steady-state noise in the cytoplasmic mRNA population counts, as quantified by the Fano Factor *FF* (the variance dived by the mean) is given by

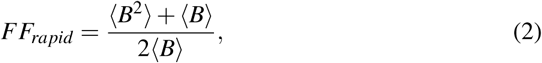

where ⟨ ⟩ is the expected value operation [29].

## 3 Results

### 3.1 General Formula Describing Noise in Gene Expression

In the section, we present a general result for the extent of fluctuations in mRNA population counts assuming the time spent by an individual RNA in the nucleus is an independent and identically distributed random variable following an arbitrary probability distribution *h*. Using the Partitioning of Poisson arrivals approach developed in [30], we obtain the following expression for the Fano factor at time *t*

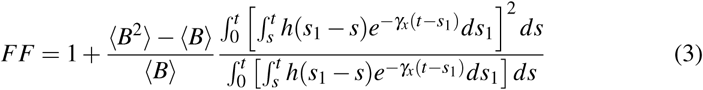

starting with zero number of RNAs in the system at *t* = 0 (details of the proof are in Appendix A). Note that in the limit of non-bursty RNA production (*B* = 1 with probability one) one obtains a Poisson distribution of mRNA counts with *FF* = 1.

We can use the general result (3) to explore the effects of diverse nuclear retention time distributions. First, we consider a delta distribution where each RNA spends a fixed time in the nucleus. Replacing *h*(*s*_1_ − *s*) in (3) with *δ* (*s*_1_ − *s*) and further simplifications results in the exact same noise

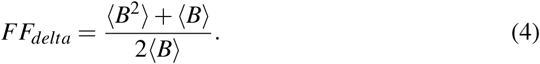

as in the case of fast RNA export (2).

Next considering a exponentially-distributed nuclear retention time with mean 1*/t*_*x*_ we substitute 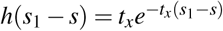 in (3) to obtain

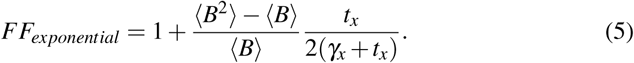

Here the noise level *FF*_*exponential*_ *< FF*_*delta*_ is smaller as compared to the delta-distributed case (4) with desynchronized RNA export events leading to effective noise buffering, As expected, (5) reduces to (2) in the limit *t*_*x*_ → ∞. Furthermore, *FF*_*exponential*_ → 1 as *t*_*x*_ → 0 which is consistent with prior experimental/theoretical findings that slow nuclear export attenuate noise in mRNA levels [31–36].

### 3.2 Extrinsic Fluctuations in Nuclear Export

Next, we consider some form of extrinsic fluctuations in the export process by considering an extrinsic factor *Z*. The stochastic dynamics of this factor is modeled analogously to the RNA production earlier, where the factor is synthesized at a Poisson rate *k*_*z*_ in random burst of size *B*_*z*_. The factor decays with rate *γ*_*z*_ that is related to the timescale of fluctuations in *Z* levels. Using (2), the steady-state coefficient of variation *CV*_*Z*_ in the level of *Z* is given by

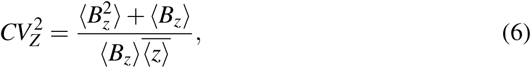

where 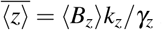 is the steady-state mean level of the extrinsic factor.

#### 3.2.1 Extrinsic Fluctuations in One-Step Delay

Extrinsic fluctuations are first incorporated into the one-step transport process, as described in Fig. 1, where the RNA export rate is now proportional to the level of *Z*. For a fixed level *z*(*t*) of *Z*, the nuclear retention time is exponentially-distributed with mean 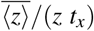. Thus, if *Z* happens to be higher than average at a given time point, then all nuclear RNA with be exported at a much faster rate. RNA export slows down as fluctuations in *Z* revert back to average levels. The overall stochastic model is described in Table 1 with stochastic events that “fire” with some probabilities in the next infinitesimal time step, and when they occur, populations counts are reset by integer amounts.

**Table 1:** Stochastic gene expression model of a one-step nuclear export process. Here *x*_0_(*t*), *x*_1_(*t*) and *z*(*t*) are integer-valued stochastic processes representing the level of the nuclear RNA, cytoplasmic mRNA and extrinsic factor *Z* at time *t*, respectively. The events describing the production of *Z* in bursts and its decay are described under the the dashed line.

| Event | Change in Population Count | Probability Event will Occur in $(t, t + dt)$ |
| --- | --- | --- |
| Transcriptional Bursting | $x_0(t) \rightarrow x_0(t) + j$ | $k_x \alpha_j dt, j = 1, 2, \dots$ |
| Nuclear Export | $x_0(t) \rightarrow x_0(t) - 1$<br>$x_1(t) \rightarrow x_1(t) + 1$ | $\frac{z(t)t_x}{\langle z \rangle} x_0(t) dt$ |
| mRNA Degradation | $x_1(t) \rightarrow x_1(t) - 1$ | $\gamma_x x_1(t) dt$ |
| ..... | ..... | ..... |
| Extrinsic Noise Production | $z(t) \rightarrow z(t) + i$ | $k_z \alpha_i dt, i = 1, 2, \dots$ |
| Extrinsic Noise Degradation | $z(t) \rightarrow z(t) - 1$ | $\gamma_z z(t) dt$ |

**Figure 1:**
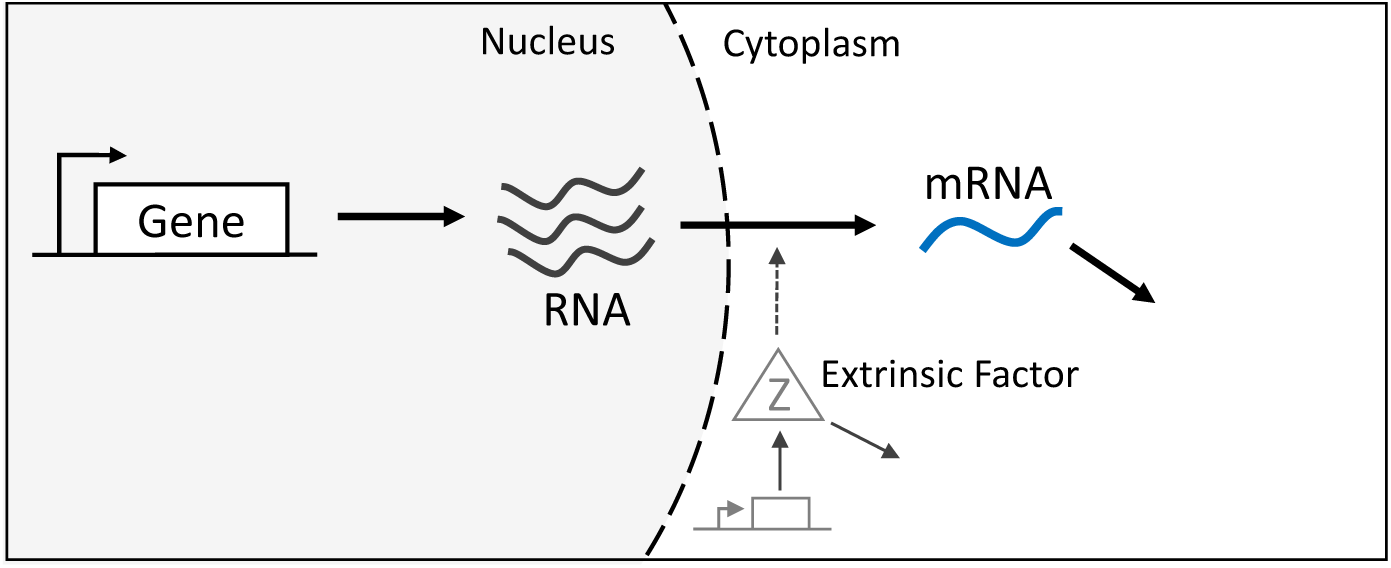
Gene expression model of the nuclear export process in the presence of an extrinsic disturbance. Within the cell nucleus, the gene synthesizes bursts of RNA transcripts that are subsequently exported to the cytoplasm to create mRNA. The cytoplasmic mRNA then decays. An extrinsic factor *Z* is modeled as a bursty birth-death process and affects the nuclear export process.

We use the Method of Moments (as explained in Appendix B) to obtain the following expression for the steady-state Fano factor of mRNA levels

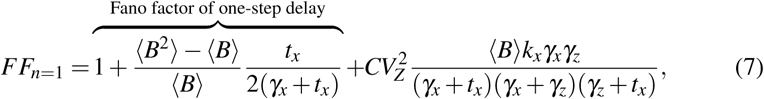

where the first term is as derived in (5) in the absence of *Z*, and the second term is the contributions from the extrinsic disturbance.

The Fano factor reveals how an extrinsic disturbance in the export process would affect downstream noise in cytoplasmic mRNA. We first consider an extremely stable extrinsic noise factor *Z* by taking the limit of *γ*_*z*_ → 0

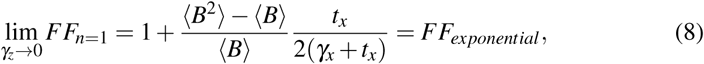

and the Fano factor simplifies to (5) - the Fano factor in the absence of any extrinsic factor. Thus, static fluctuations in *Z* are not transmitted downstream to the mRNA level. We next consider how average RNA transport time affects mRNA noise, and find that the mRNA Fano factor is not always buffered by increasing RNA transport time as before. Results in Fig. 2 depict noise buffering at increased RNA transport time when the extrinsic factor is not incorporated. However, when the extrinsic factor is included, the mRNA Fano factor exhibits non-monotonic behavior, where the Fano factor initially decreases before increasing, in relation to the average RNA transport time. The extent of the increased Fano factor at higher average RNA transport times is additionally influenced by the variability and the mean level of the extrinsic factor. This indicates that in the presence of extrinsic fluctuations, short average RNA transport time is ideal as it offers better mRNA noise buffering, while increased RNA transport time increases mRNA noise.

**Figure 2:**
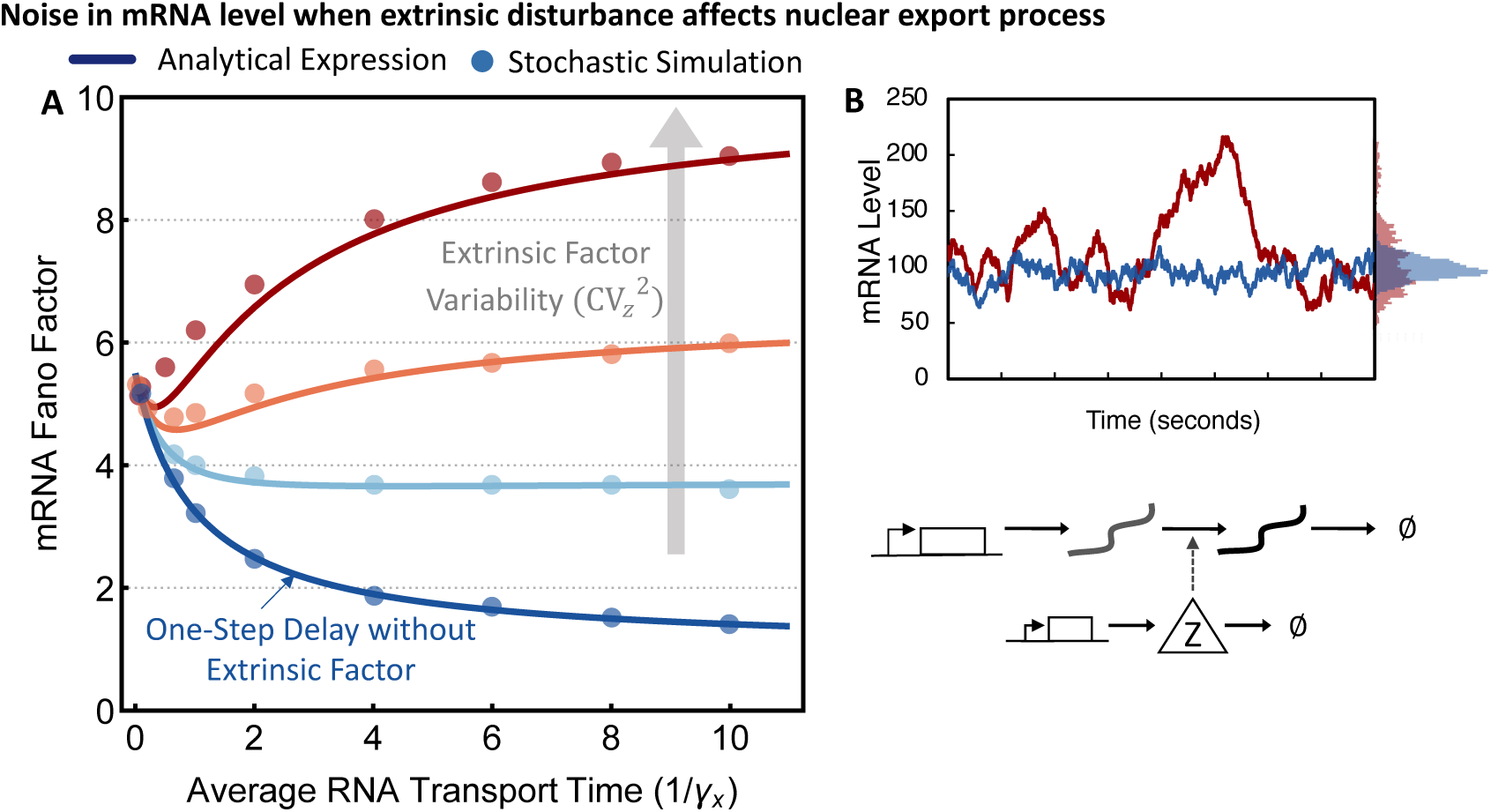
**(A)** Noise in mRNA level for a one-step transport process is plotted as a function of increasing transport time, normalized by the mRNA decay rate when extrinsic disturbances affect the transport process. The extrinsic factor, modeled by species *Z*, is altered to become more or less noisy. The dark blue line denotes the mRNA Fano factor without the extrinsic disturbance. The effect of increasing variability in extrinsic factor level, quantified by the extrinsic factor level coefficient of variation squared, is depicted by the light blue, orange, and red lines, with the values of 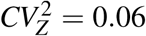, 0.1, and 0.6. Monte Carlo Simulations (SSA) results are denoted by circle markers and validate the analytical formulas. **(B)** Example traces of mRNA level fluctuations with time. The red line denotes mRNA level in the presence of an extrinsic factor with 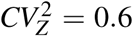 while the blue line denotes mRNA level in the presence of an extrinsic factor with 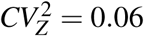. The remaining parameters have the following values: ⟨*B*⟩ = 10, *k*_*x*_ = 10, ⟨*B*_*z*_⟩ = 10, *γ*_*z*_ = 1, *t*_*x*_ = 1.

We further explore the extent of the extrinsic factor dynamics on shaping mRNA noise by considering the extrinsic species time scale. We again find interesting non-monotonic behavior in the mRNA Fano factor, depicted in Fig. 3, where noise is plotted as a function of the extrinsic factor decay rate (*γ*_*z*_). The mRNA Fano factor increases at low extrinsic factor time scales. However, as the timescale is increased, the mRNA noise is buffered. We additionally find that when the extrinsic factor has increased variability, the mRNA Fano factor is also increased.

**Figure 3:**
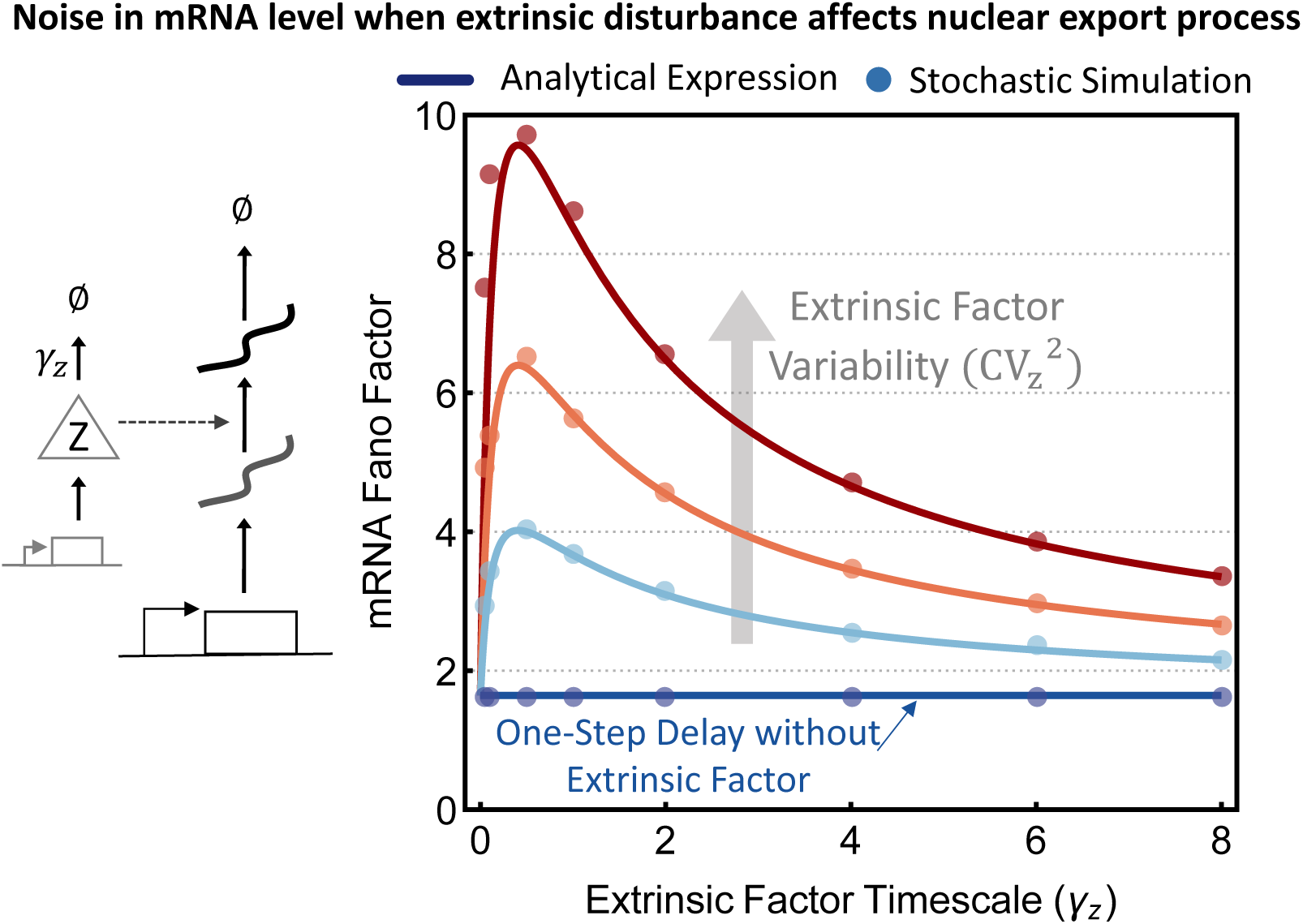
Cytoplasmic mRNA level noise as a function of extrinsic factor timescale. For each case, the mean level of extrinsic factor is held constant. mRNA Fano factor is plotted for increasing variation in the extrinsic factor level, quantified by its coefficient of variation squared, taking on the following values: 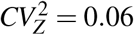, 0.1, and 0.6 denoted by the light blue, orange, and red slopes, accordingly. The dark blue line depicts the mRNA Fano factor when an extrinsic factor is *not* included. Monte Carlo Simulations (SSA) results are denoted by circle markers and validate the analytical formulas. The remaining parameters have the following values: ⟨*B*⟩ = 10, *k*_*x*_ = 10, ⟨*B*_*z*_⟩ = 10, *γ*_*z*_ = 1, *t*_*x*_ = 1.

#### 3.2.2 Extrinsic Fluctuations for a Multi-Step Time Delayed Export

Extrinsic disturbances are next incorporated into a time delayed nuclear export process. Here, the export process is modeled as multiple events, implemented through irreversible conversion reactions converting the nuclear RNA *X*_0_ into cytoplasmic mRNA *X*_*n*_ by intermediate states: *X*_1_, …*X*_*n*−1_, where the rate of conversion is given by 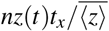. Note that for a fixed *z*, this corresponds to an Erlang-distributed nuclear retention time. For a two-step export process the overall stochastic model is illustrated in Table II

**Table 2:** Stochastic gene expression model of a two-step nuclear export process (*n* = 2; the incorporation of extrinsic noise species *Z* is described under the the dashed line. Here *x*_*i*_(*t*) denotes the population count of species *X*_*i*_ at time *t*.

| Event | Change in Population Count | Probability Event will Occur in $(t, t + dt)$ |
| --- | --- | --- |
| Transcriptional Bursting | $x_0(t) \rightarrow x_0(t) + j$ | $k_x \alpha_j dt, j = 1, 2, \dots$ |
| Nuclear Export | $x_0(t) \rightarrow x_0(t) - 1$<br>$x_1(t) \rightarrow x_1(t) + 1$ | $\frac{2t_x z(t)}{\langle z \rangle} x_0(t) dt$ |
| Nuclear Export | $x_1(t) \rightarrow x_1(t) - 1$<br>$x_2(t) \rightarrow x_2(t) + 1$ | $\frac{2t_x z(t)}{\langle z \rangle} x_1(t) dt$ |
| mRNA Degradation | $x_2(t) \rightarrow x_2(t) - 1$ | $\gamma_x x_2(t) dt$ |
| ..... | ..... | ..... |
| Extrinsic Noise Production | $z(t) \rightarrow z(t) + i$ | $k_z \alpha_i dt, i = 1, 2, \dots$ |
| Extrinsic Noise Degradation | $z(t) \rightarrow z(t) - 1$ | $\gamma_z z(t) dt$ |

Using the Method of Moments, we derive the following formula for the steady-state mRNA Fano factor corresponding to a two-step time delay (*n* = 2) in the presence of an extrinsic disturbance

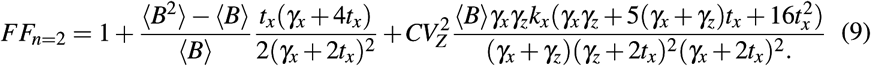

Note by taking 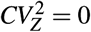 one would obtain the same Fano factor as obtained using (3) for an Erlang-distributed nuclear retention time.

Using a similar process the Fano factor can be derived for a three-step time-delayed export process (*n* = 3). Results for the one-step and three-step transport process are compared in Fig. 4. The Fano factor is plotted as a function of increasing average RNA transport time and is shown for when the extrinsic factor is, and is not included. First, we note that when the one-step and three-step transport processes are not in the presence of extrinsic fluctuations, the mRNA noise is similar. The mRNA noise becomes the same at increased average RNA transport time. Next, we examine the one-step and three-step transport processes when the extrinsic factor is included. The mRNA Fano factor exhibits greater noise when extrinsic fluctuations are included in the three-step process compared to the one-step process. Additionally, when nuclear export has a one-step or three-step delay, the Fano factor exhibits non-monotonic behavior, where the non-monotonic behavior is more apparent in the three-step delay. Last, when average RNA transport time is increased, the Fano factor for the one-step and three-step delay eventually results in the same mRNA Fano factor.

**Figure 4:**
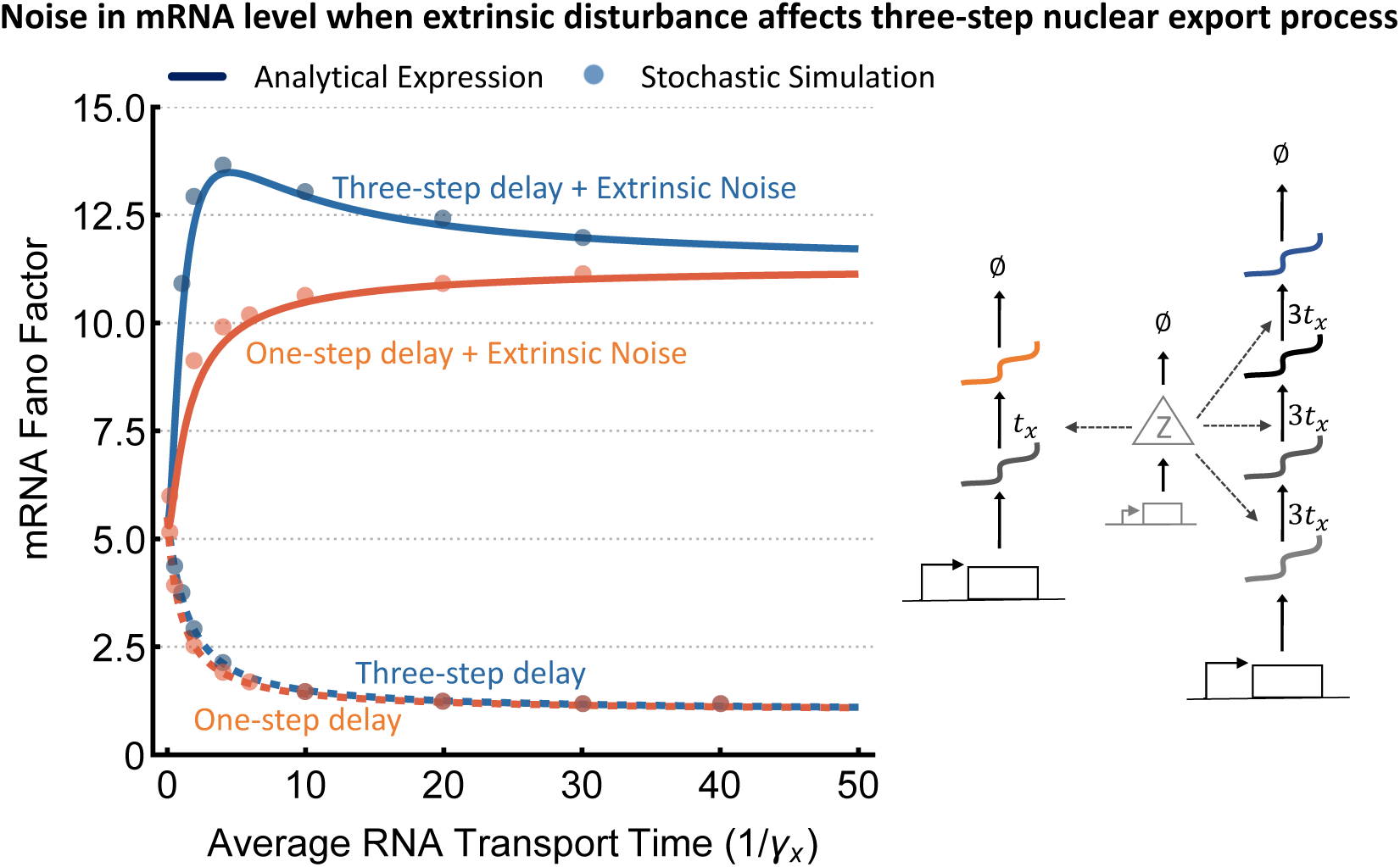
Cytoplasmic mRNA level noise plotted as a function of increasing nuclear RNA transport time, normalized by the mRNA decay rate, comparing time delayed nuclear export with and without the presence of extrinsic noise, denoted by solid and dashed lines, accordingly. The three-step export process is denoted by blue and the one-step delay is denoted by orange. The analytical formulas (lines) are validated by Monte Carlo Simulations (SSA), denoted by the circle markers. The remaining parameters have the following values: ⟨*B*⟩ = 10, *k*_*x*_ = 10, *k*_*z*_ = 4, and *γ*_*z*_ = 3.

## 4 Conclusion

We have systematically studied the role that different characteristics of the nuclear export process play in shaping the extent of random fluctuations in the mRNA level of a given gene. We considered how mRNA noise is affected by different nuclear retention time distributions, by time-delayed nuclear export, and by extrinsic fluctuations in nuclear export. In regards to nuclear export timing, a general observation found is that increased average export time, thus a small export rate buffers cytoplasmic mRNA noise.

Specifically, there are regimes where increased variability in RNA transport time actually buffers mRNA noise. Additionally, when extrinsic fluctuations are incorporated, mRNA noise exhibits non-monotonic behavior when plotted as a function of extrinsic species timescale, where noise is originally low for stable extrinsic species, then sharply increases as extrinsic species stability is decreased, before decreasing again for extremely unstable extrinsic species. Furthermore, the effects of extrinsic fluctuations are magnified when the fluctuations are incorporated into *n*−step delayed nuclear export.

## A Fano Factor using Partitioning of Poisson Arrivals Method

We apply the mapping based on partitioning of Poisson arrivals (PPA-mapping) [30] to consider a reduced model with bursts arriving at rate *k*_*x*_*/N* where *N* → ∞. There-fore, the mRNA generating functions for the the original *G*(*y, t*) and the reduced *g*(*y, t*) models are related by

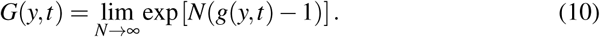

In the next step we derive *g*(*y, t*). For the reduced model, bursts are arriving with rate *k*_*x*_*/N*, i.e., the number of burst arrivals at time *t* is a Poisson-distributed random variable with mean *k*_*x*_*t/N*. In the limit of *N* → ∞, we only need to consider the 0-burst and 1-burst arrival cases in time *t*. Correspondingly, we have

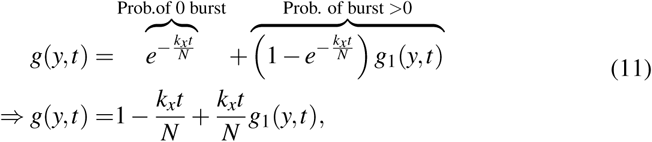

where *g*_1_(*y, t*) is the generating function of mRNA distribution at time *t*, conditional on exactly 1 burst arrival between time 0 to *t*. To this end, in order to calculate *G*(*y, t*) we need to calculate *g*_1_(*y, t*).

Consider a single mRNA is created at *t* = *s*. Conditioning on the export time (*s*_1_), the generating function for the distribution of mRNAs at time *t* is given by

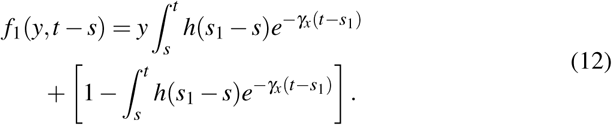

Let *g*_*b*_(*y*) be the generating function for probability distribution of number of RNAs produced in the burst. Then, by conditioning on the time of burst arrival *t* = *s* we have

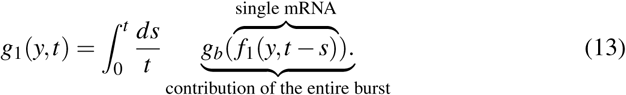

The above expression is conditioned on having a single bursting event which means the burst arrival time distribution is uniform over [0, *t*]. Substituting (13) in (11), and then using (10) we get

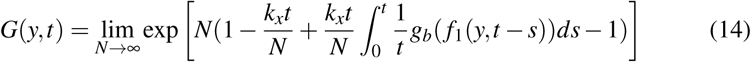

simplifying to

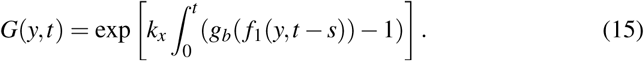

By having the generating function we can calculate the mean and variance of the mRNA counts *x*_1_(*t*) by using the following formulas

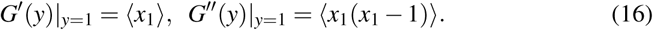

Hence, we can derive the mean count as

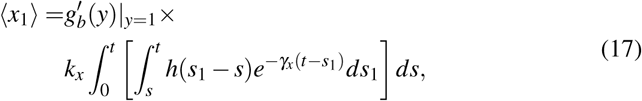

and its second-order moment as

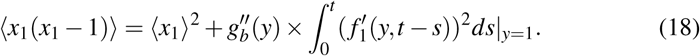

By replacing the burst size’s first- and second-order moments as

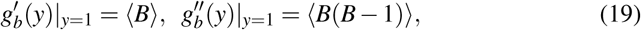

we have

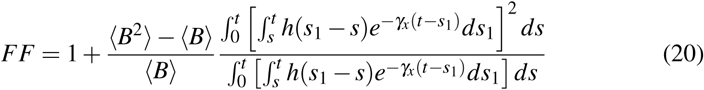

Let *τ* denote the mean waiting time for mRNA decay including RNA export time. Then by using Little’s law we have

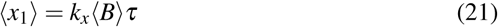

Hence from (17) we have

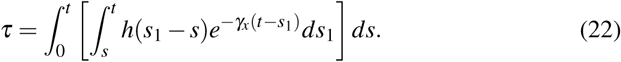

## B Fano Factor using Method of Moments

To characterize the effects of extrinsic noise in the nuclear export rate, we again obtain the Fano factor. To include the extrinsic noise species, *Z*, we utilize the Method of Moments to obtain the Fano factor of the export model described by the stochastic formulation described in Table 1. In this table, the extrinsic noise factor is described in the rows below the dashed line. We assume each event is probabilistic and occurring at exponentially-distributed time intervals. When an event occurs, the nuclear RNA and cytoplasmic mRNA population count must be updated, as described by the resets in the center column of the table. The propensity functions in the rightmost column describe the probability that the event will occur in the next infinitesimal time. The time derivative of the expected value of any differentiable function *ϕ*(*x*_0_, *x*_1_, *z*) is given by [37]:

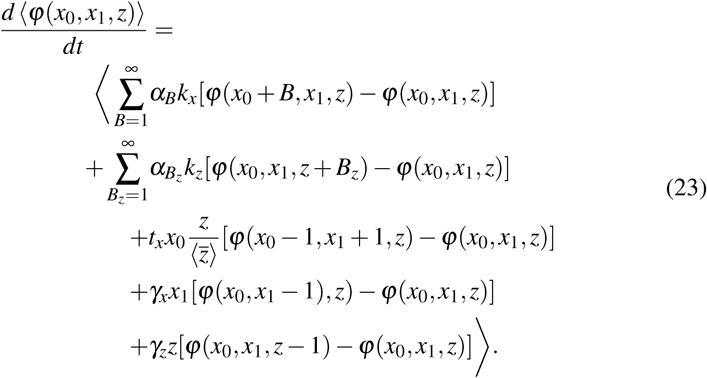

The Fano factor is found by first selecting appropriate choices for the function *ϕ*(*x*_0_, *x*_1_, *z*) to obtain the first- and second-order moments [37]. We then solve for steady-state to determine the mean and variance of mRNA level. In this model, non-linearity is introduced in the export rate which takes the form *t*_*x*_*zx*_0_. The non-linearity causes the system to become analytically intractable. To obtain the Fano factor we linearize this rate using Taylor Series expansion. We assume small fluctuations in nuclear RNA count *x*_0_(*t*) and extrinsic noise factor count *z*(*t*) [38–40]. By linearizing the transport rate about its respective means 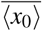 and 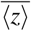, we approximate the following transport rate as

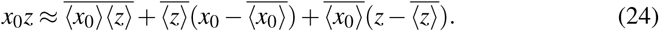

When obtaining the moment dynamics, we replace the non-linear transport rate in (23) with the linear rate (24).

Selecting appropriate choices for *ϕ*(*x*_0_, *x*_1_, *z*) and using the resets and propensity functions described in Table I, we reveal the following moment dynamics:

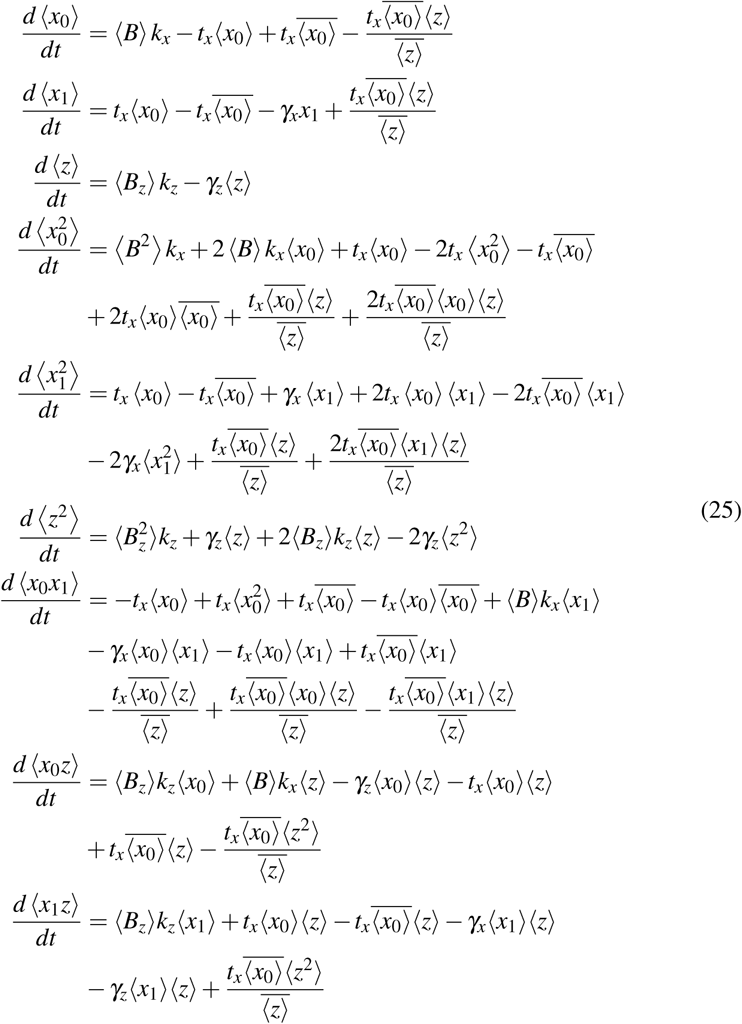

[41, 42]. Quantifying the steady-state first- and second-order moments results in the Fano factor (7).

## ACKNOWLEDGMENT

This work is supported by the NSF grant ECCS-1711548 and ARO grant W911NF1910243 to AS, and NSF grant DMS-1854350 to RVK.

